# Structural Basis of *Streptomyces* Antibiotic Regulatory Proteins Activating Transcription

**DOI:** 10.1101/2023.09.04.556155

**Authors:** Yiqun Wang, Xu Yang, Feng Yu, Zixin Deng, Shuangjun Lin, Jianting Zheng

## Abstract

Streptomycetes are renowned antibiotic producers, with *Streptomyces* antibiotic regulatory proteins (SARPs) acting as activators for antibiotic biosynthesis. However, the precise mechanism underlying SARPs’ transcriptional activation remains elusive. Here, we used cryo-electron microscopy (cryo-EM) to unravel the interplay between SARP, DNA, and RNA polymerase (RNAP) during transcriptional activation. The SARP domain of *Streptomyces coelicolor* AfsR (SAS) forms a side-by-side dimer contacting the *afs box* centered at −29.5 relative to the transcription start site. The upstream protomer binds to the direct repeat encompassing the −35 element while the σ^HrdB^ region 4 (R4) is positioned on top of both protomers, causing the removal of R4 from the major groove of the −35 element. Both SAS protomers establish interactions with C-terminal domain of one RNAP α subunits, while specific regions of the RNAP β flap tip helix and β’ zinc-binding domain also engage with SAS. Key interfacial residues accounting for transcriptional activation were confirmed by mutational studies and *in vitro* transcriptional assays. Overall, our results present a detailed molecular view of how SARPs serve to activate transcription.

## 1. Introduction

Streptomycetes are multicellular bacteria with a complex developmental cycle(*1*). They produce numerous bioactive natural products, including many antibiotics with important applications in medicine and agriculture. Genome analyses of streptomycetes in recent years have revealed their substantial biosynthetic potential. This capability makes streptomycetes one of the most important industrial microbial genera. Complex regulatory systems tightly govern the natural product biosynthesis of streptomycetes, involving both global and pathway-specific regulators. *Streptomyces* antibiotic regulatory protein (SARP) family regulators are prominent initiators and activators of natural product biosynthesis, and have mainly been found in actinomycetes(*2*). Recently reported members of the SARP family include ChlF2 from *Streptomyces antibioticus* and MilR3 from *Streptomyces bingchenggensis*. ChlF2 activates the production of chlorothricin, known for its anti-inflammatory properties, and MilR3 stimulates the synthesis of an excellent insecticide milbemycin (*3, 4*). SARP regulators exhibit a comparable protein architecture and binding affinity towards similar recognition sequences (*5*). These regulators feature an N-terminal OmpR-type DNA-binding (ODB) domain and a C-terminal bacterial transcriptional activation (BTA) domain, collectively referred to as the SARP domain. The ODB domain is characterized by a winged helix-turn-helix (HTH) comprising three α-helices and two antiparallel β-sheets(*6*), whereas the BTA domain is all-helical(*7*). In contrast to “small” SARPs that only contain a SARP domain, “large” SARPs possess extra domains at their C-terminals that are hypothesized to modulate the activity of the SARP domain (Fig. 1A). “Small” SARPs are more frequently observed and extensively studied among actinomycetes, compared to their “large” counterparts (*8*).

**Fig. 1.**
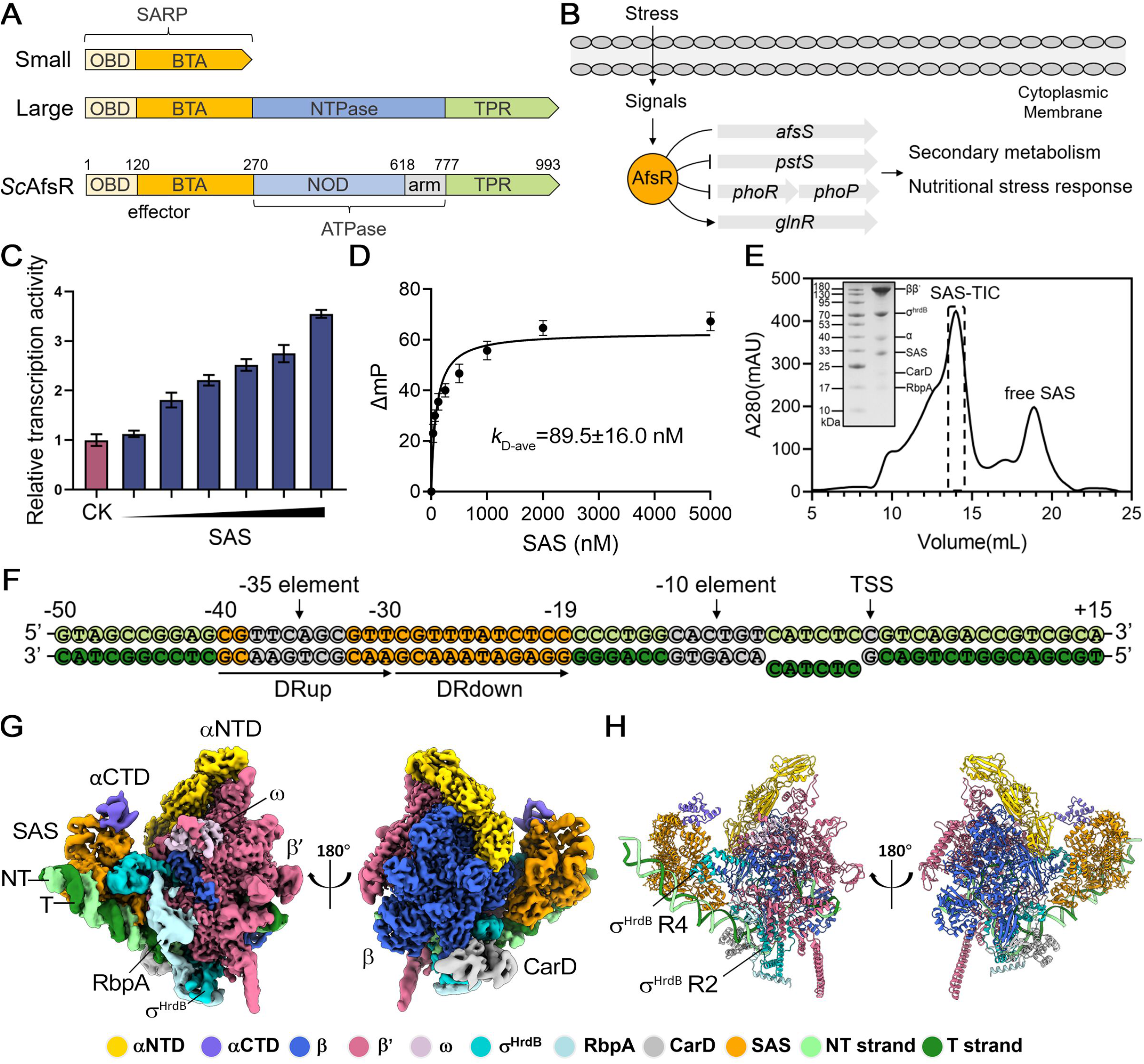
The overall structure of SAS-TIC. (A) Domain structures of *Streptomyces* antibiotic regulatory protein (SARP) regulators and *Streptomyces coelicolor* AfsR (*Sc*AfsR). The N-terminal ODB (OmpR-type DNA-binding) domain together with the following BTA (bacterial transcriptional activation) domain is referred to a SARP domain. TPR, tetratricopeptide repeat; NOD, nucleotide-binding oligomerization domain. (B) Hypothetical scheme for the regulation by AfsR in *Streptomyces coelicolor*. (C) Transcription assays with increasing concentrations (62.5 nM, 125 nM, 250 nM, 500 nM, 750 nM, 1000 nM) of SAS. Data are presented as mean ± SEM from three independent assays. (D) Fluorescence polarization assay of SAS with *afs box*. The concentration of *afs box* was 10 nM. Error bars represent mean ± SEM of n=3 experiments. (E) Assembly of SAS-transcription initiation complex (TIC). The protein compositions in the dotted line boxed fractions are shown in the SDS-PAGE. (F) The *afsS* promoter fragment used for cryo-EM structure determination. The -35 element, -10 element, the transcription start site (TSS) and the 6-bp noncomplementary bubble are denoted. The *afs box* is colored orange and contains DRup (the upstream direct repeat) and DRdown (the downstream direct repeat). The top (non-template) strand and bottom (template) strand are colored light green and dark green, respectively. (G) Two views of cryo-EM map. The map was generated by merging the consensus map of the full SAS-TIC and the focused map of the αCTD-SAS-DNA in Chimera X. NT, non-template; T, template. (H) Cartoon representation structure of SAS-TIC. The RNAP subunits, DNA, and regulators are colored as in the color scheme.

*Streptomyces coelicolor* AfsR (*Sc*AfsR) is a pleiotropic, global regulator of secondary metabolism belonging to the “large” SARP group, containing a SARP (Met1-Ala270) domain, a conserved ∼35-kDa dubbed nucleotide-binding oligomerization domain (NOD) (Ala271 to Glu618), an arm domain (Arg619-Glu777) as well as a tetratrico-peptide repeats (TPR) sensor domain (Asp778-Arg993) (*9*) (Fig. 1A). It activates *afsS* transcription by binding to the promoter(*10, 11*). AfsS is a master regulator of both secondary metabolism and nutritional stress response(*12*). It was presumed that upon binding of AfsR to a 22-base pair (bp) binding box (*afs box*), RNA polymerase (RNAP) holoenzyme is recruited to the *afsS* promoter and forms a ternary DNA-AfsR-RNAP complex, allowing for transcriptional initiation(*9*). Additionally, *Sc*AfsR also binds to promoters of PhoR-PhoP regulon members such as *pstS, phoRP* and *glnR*, controlling the response to phosphate and nitrogen scarcity (*13, 14*) (Fig. 1B). *In vitro*, a truncated *Sc*AfsR protein containing only the SARP domain (SAS) recognizes the *afsS* promoter and activates its transcription as the full-length AfsR does, suggesting that the DNA-binding specificity of AfsR and its capability to activate *afsS* promoter are determined solely by SAS (*9*). However, the detailed mechanism mediating SAS-dependent transcription activation is still obscure.

The RNAP holoenzyme consists of a dissociable σ subunit that establishes specific interactions with the −35 and −10 elements of the promoter in order to initiate transcription. The positioning of RNAP is primarily influenced by the contacts between σ region 4 (R4) and the −35 element, while the formation of an open complex is driven by the contacts between σ region 2 (R2) and the −10 element(*15, 16*). The *afsS* promoter comprises a consensus −10 element (CACTGT) but a suboptimal −35 element (TTCAGC). This characteristic is commonly observed in streptomycete promoters (*17, 18*), which complicates the identification of the −35 elements. The 22-bp *afs box*, centered at position −29.5 relative to the transcription start site in the *afsS* promoter, overlaps with the spacer between the −10 and −35 elements. It is composed of two 11-bp direct repeats (DRs), with the upstream DR (−40 to −30) overlapping with the −35 element (−38 to −33)(*19, 20*). The catabolite activator protein (CAP), also known as the cAMP receptor protein (CRP), is an extensively studied transcription factor. The 22-bp CAP binding box (*cap box*) is centered at –61.5 in class I promoters and at –41.5 in class II promoters(*21-23*). The binding position of SAS suggests that it may possess a distinct activation mechanism.

Here, we used cryo-electron microscopy (cryo-EM) to visualize a SAS-transcription initiation complex (TIC) and to provide a transcription-activating mechanism that could be generally used by SARP family activators.

## 2. Results

### Overall structure of SAS-TIC

*Sc*AfsR, a prominent global regulator of secondary metabolism, possesses a widely distributed SARP domain in all actinomycetes, enabling the activation of antibiotic synthesis gene clusters. This characteristic has sparked interest in the function of *Sc*AfsR, with particular emphasis on its SARP domain, namely SAS. *In vitro* MangoIII-based transcription assays demonstrated that SAS enhances transcription of *afsS* promoter (Fig. 1C), in agreement with the previous report (*9*). A 32 bp DNA probe (−45 to −14) containing the *afs box* (−19 to −40) was used for the fluorescence polarization assays of SAS, in which an average dissociation constant *k*_D-ave_ of 89.5 ± 16.0 nM was obtained (Fig. 1D). SAS purified to homogeneity was directly used to assemble the activator-TIC (Fig. S1). A DNA scaffold is engineered from the *afsS* promoter, consisting of a 44-bp (−50 to −7) upstream promoter double-stranded DNA (dsDNA) which contains the 22-bp *afs box* and the consensus –10 element, a 6-bp (−6 to −1) non-complementary transcription bubble, and a 15-bp (+1 to +15) downstream promoter dsDNA (Fig. 1F). RbpA and CarD, two RNAP-binding proteins discovered in actinobacteria, can stabilize the TIC and have been found in many TIC structures of *Mycobacterium tuberculosis (24-26*). Therefore, we combined SAS, the DNA scaffold, RbpA, CarD, and RNAP σ^HrdB^-holoenzyme, and subsequently separated the TIC complex using size exclusion chromatography (Fig. 1E). Through *in vitro* transcription assays, we observed that the presence of CarD and RbpA during the transcription of the *afsS* promoter yielded results almost the same as those achieved with the RNAP holo-enzyme alone. Furthermore, SAS maintained its role as a transcriptional activator in the presence of RbpA and CarD (Fig. S2).

The final cryo-EM map of the SAS-TIC was reconstructed using a total of 95223 single particles and refined to a nominal resolution of 3.35 Å, with ∼3 Å at the center of RNAP and ∼6 Å at the peripheral SAS (Fig. S3). Local refinement focused on the SAS region generated a 3.77-Å-resolution map. In the cryo-EM maps, the densities allowed unambiguous docking of two SAS protomers, a 61-bp promoter DNA (−46 to +15), a σ^HrdB^ subunit except disordered region 1.1, five subunits of RNAP core (α_2_ββ′ω), RbpA and CarD. In addition, the cryo-EM density map shows a clear signal for an αCTD (Fig. 1G). The two SAS protomers simultaneously engage the *afs box* and RNAP σ^HrdB^-holoenzyme. The overall structure of *S. coelicolor* RNAP in SAS-TIC closely resembles that in our recently reported *S. coelicolor* RNAPσ^HrdB^-Zur-DNA structure (PDB ID: 7X75, rmsd of 0.334 Å for 3002 aligned Cα atoms)(*27*), and other actinobacteria RNAP structures including *M. tuberculosis* RNAP-promoter open complex (PDB ID: 6VVY, rmsd of 1.133 Å for 2745 aligned Cα atoms)(*28*) and *Mycobacterium smegmatis* RNAP TIC (PDB ID: 5VI5, rmsd of 1.048 Å for 2621 aligned Cα atoms)(*29*) (Fig. S4).As shown in Fig. 1H, σ^HrdB^ R4 is positioned on top of two SAS protomers, instead of binding in the major groove of the −35 element as observed in the reported structures (*27-29*), suggesting a sigma adaptation mechanism like *M. smegmatis* PafBC (*30*). CarD binds the β-subunit and interacts with the upstream double-stranded/single-stranded (ds/ss) junction of the transcription bubble (*31*), while RbpA enhances interactions between β’, σ^HrdB^ and promoter DNA(*32*).

### Interactions between SAS and *afs box*

The SAS-TIC structure elucidates the binding interactions between SAS protomers and two major grooves of the *afs box*, with each protomer making contacts with one DR (Fig. 2A). The contacts span the first eight conserved base pairs. In contrast, the last 3 base pairs don’t make any contact with SAS (Fig. 2B). Overall conformations of the two protomers are essentially the same with an overall rmsd of 0.4 Å. A dimer interface of ∼1120 Å^2^ is formed between the two SAS protomers, revealing a unique side-by-side arrangement (Fig. 2C). Consistent with the previous report (*9*), size exclusion chromatography showed that SAS is monomeric in solution (Fig. S1B). This suggests that the dimerization of SAS is occurred specifically upon DNA binding. As a representative example of SARP family transcription activators, the SAS folds into an N-terminal ODB domain (residues 25-111) and a C-terminal BTA domain (residues 120-270) (Fig. 2D). The ODB domain consists of three α helices packed against two antiparallel β sheets. Two helices, α2 and α3, and the seven-residue connecting loop constitute an HTH DNA-binding motif, followed by a β hairpin that contacts the strand between the helices α1 and α2. The BTA domain comprises seven α-helices, of which the first three ones stand on the first β sheet and the first two helices of the N-terminal ODB domain, burying an interfacial area of ∼860 Å^2^. The overall structure of SAS protomers closely resembles EmbR (PDB ID: 2FEZ, rmsd of 2.8 Å for 247 aligned Cα atoms) (*7*), a transcriptional regulator of *M. tuberculosis*, except that EmbR comprises an additional C-terminal forkhead-associated (FHA) domain (Fig. S5).

**Fig. 2.**
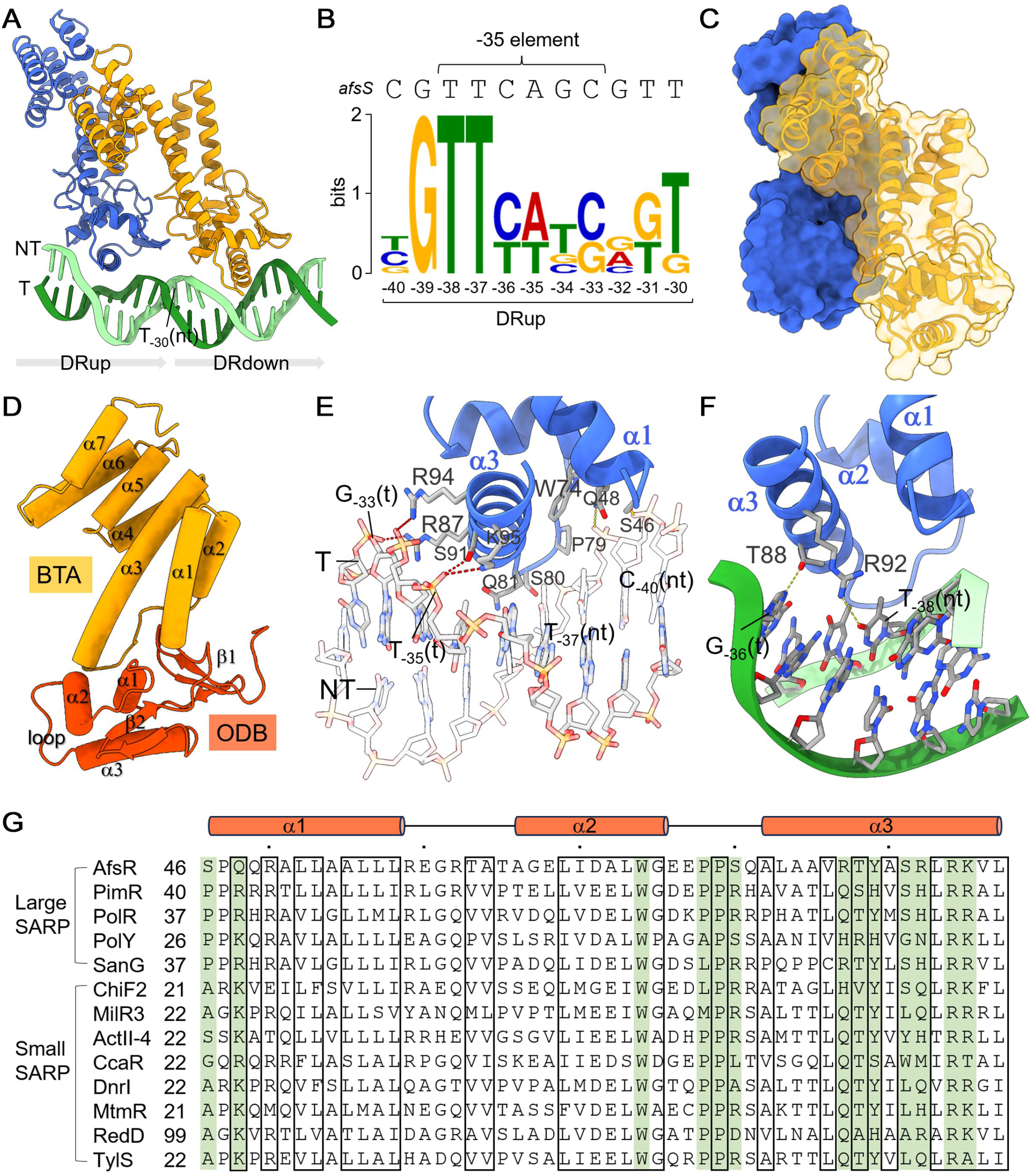
Interactions of SAS with *afs box*. (A)Two SAS protomers bind to the *afs box* (-19 to -40) side-by-side with each protomer contacting one DR. T_-30_ (nt) is the last nucleotide of DRup. Upstream and downstream SAS are colored blue and orange, respectively. (B) Conserved sequences corresponding to the 11-nt DR of the AfsR operators generated by MEME. (C) The dimer interface between two SAS protomers. (D) The domain organization indicated in the downstream SAS protomer. ODB, OmpR-type DNA-binding; BTA, bacterial transcriptional activation. (E) Detailed interactions of upstream SAS with the DNA backbone phosphate groups. The R87, S91, R94 and K95 in helix α3 form salt bridges with T_-35_ to G_-33_ of the t-strand. The N-terminal end of helix α1(S46 and Q48), the C-terminal end of helix α2 (W74), and the HTH loop (P79, S80, Q81) make hydrogen bonds and van der Waals interactions with C-40 to T_-37_ of the nt-strand. Hydrogen bonds and salt bridges are shown as yellow and red dashed lines. (F) Contacts of SAS with specific nucleotides. The residues R92 and T88 make hydrogen bonds (shown as yellow dashed lines) with O4 of T_-38_(nt) and N7 of G_-36_(t) respectively. (G) Sequence alignments of SARP regulators from different *Streptomyces* strains, highlighting the residues interacting with DNA (green). Only ODB domains were compared. These proteins include AfsR (P25941), ActII-4 (P46106), RedD (P16922) from *S. coelicolor*, PimR (Q70DY8) from S. natalensis, PolY (ABX24502.1), PolR (ABX24503.1) from *S. asoensis*, SanG (Q5IW77) from *S. ansochromogenes*, ChlF2 (Q0R4N4) from *S. antibioticus*, MilR3 (D7BZQ7) from *S. bingchenggensis*, CcaR (P97060) from *S. clavuligerus*, DnrI (P25047) from *S. peucetius*, MtmR (Q194R8) from *S. argillaceus* and TlyS (M4ML56) from *S. fradiae*. The black boxes highlight the positions conserved.

The two SAS protomers establish a total contact surface of ∼1300 Å^2^ with the dsDNA, resulting in a 20° bend of the helical axis of the upstream dsDNA at T_-30_(nt) (-30T on the non-template strand) (Fig. S6). The two ODB domains make similar contacts with DNA, inserting the helix α3 of the HTH into the major groove of the DR. The DNA contacts made by the upstream SAS protomer were described in the following section (Fig. 2E). The N-terminal end of helix α1, the C-terminal end of helix α2, and the HTH loop make extensive contacts with the backbone phosphate groups from C-_40_ to T_-37_ of the nt-strand, involving both hydrogen bonds (S46 and Q48) and van der Waals interactions (W74, P79, S80 and Q81). The recognition helix α3 penetrates the DNA major groove and is almost perpendicular to the DNA axis. R87, R94 and K95 form salt bridges with the backbone phosphate groups from T_−35_ to G_-33_ of the t-strand. The side chain of S91 makes a hydrogen bond with the phosphate group of T_−35_(t). In addition to these nonspecific contacts with backbone phosphate groups, the recognition helix α3 also establishes specific DNA contacts (Fig. 2F). The R92 makes hydrogen bonds with the O4 atom of T_-38_(nt) via its guanidinium group. The side chain of T88 makes a hydrogen bond with N7 of G_-36_(t). Consistent with the structural observations, previous EMSAs show that changing the T-_38_(nt) of the upstream DR or the corresponding T-_27_(nt) of the downstream DR to adenosine prevents the binding of *Sc*AfsR (*20*). However, changing the G_-36_(t) to adenosine doesn’t impair the binding of *Sc*AfsR since A_-25_(t) is observed at the corresponding position of the downstream DR (*20*). Most residues involved in DNA interactions are highly conserved in both “small” and “large” SARP homologues across various *Streptomyces* strains (Fig. 2G), including Q48, W74, P79, T88, R94, and K95.

### SAS engages with σ^HrdB^ R4, β FTH, and β’ ZBD

Both SAS protomers make contacts with σ^HrdB^ R4, burying a total interface of 800 Å^2^ (Fig. 3A). The upstream protomer contacts σ^HrdB^ R4 by its ODB domain, of which the HTH forms a concave surface at the N-terminal of the helix α3 to enfold the N-terminal of the helix α4 of σ^HrdB^ R4. The negatively charged residues E76 and E77 in the loop connecting helices α2 and α3, and the residues E68 and D71 in helix α2 of the ODB domain form salt bridges with R485 and R487 in σ^HrdB^ R4, respectively (Fig. 3B). The interface where the shape and charge complement each other buries an area of ∼320 Å^2^. The downstream protomer mainly uses its BTA domain to contact σ^HrdB^ R4, with a buried surface area of ∼480 Å^2^ (Fig. 3C). The helices α3 to α7 of the downstream BTA domain form a “chair” to let σ^HrdB^ R4 sit on it. The C-terminal of helix α4, the N-terminal of helix α5 and the connecting turn of σ^HrdB^ R4 fit in a groove formed by helices α3 to α7 of the BTA domain. The R498 at the C-terminal of helix α4 of σ^HrdB^ R4 makes contacts with the E176 and V172 on the “chair back”. The H499 in the connecting turn of σ^HrdB^ R4 is enfolded in an amphiphilic pocket formed by residues L211, L243, L247, V249, E213, and R252 of the BTA domain. The K496 at the C-terminal of helix α4 of σ^HrdB^ R4 makes a salt bridge with the E246 at the C-terminal of helix α6 of the BTA domain. In addition, the HTH loop of the ODB domain forms interactions with the loop connecting helices α2 and α3 of σ^HrdB^ R4.

**Fig. 3.**
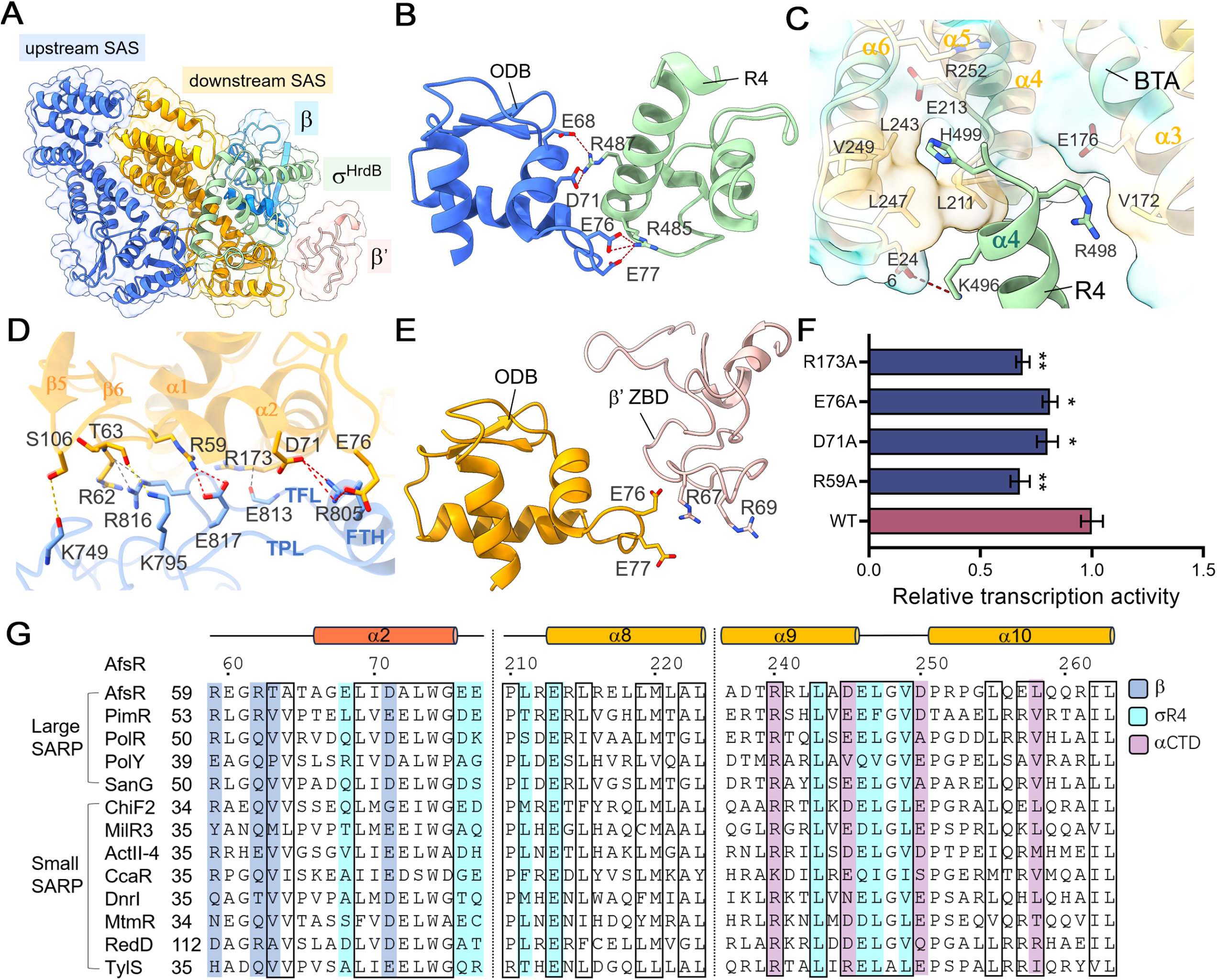
SAS interacts with σ^HrdB^ R4, β FTH, β’ ZBD. (A) The SAS protomers interact with σ^HrdB^, β and β’. (B) The upstream SAS protomer contacts σ^HrdB^ R4 by its ODB domain. The loop connecting helices α2 and α3 (E76 and E77), and helix α2 (E68 and D71) form salt bridges (shown as red dashed lines) with R485 and R487 in σ^HrdB^ R4. (C) The helices α3 to α7 of the downstream BTA domain form a “chair” to let σ^HrdB^R4 sit on it. The H499 in the connecting turn of σ^HrdB^R4 is enfolded in an amphiphilic pocket of the BTA domain as shown. The K496 at the C-terminal of helix α4 of σ^HrdB^R4 makes a salt bridge with the E246 of the BTA domain. (D) Both ODB and BTA domains of the downstream SAS protomer make extensive interactions with the β FTH, the preceding loop (TPL) and the following loop (TFL) of the β flap. SAS is colored orange and β flap is colored blue. Hydrogen bonds, salt-bridges and van der Waals interactions are shown as yellow, red and grey dashed lines, respectively. (E) Interactions between the β′ ZBD and the ODB of downstream SAS. The positively charged R67 and R69 of β′ ZBD contact the negatively charged E76 and E77 of the HTH loop of the ODB domain. (F) Mutating interfacial residues of SAS impaired transcription activation. Data are presented as mean ± SEM from three independent assays. ^*^ *P* < 0.05; ^**^ *P* < 0.01 in comparison with the wild-type SAS. (G) Sequence alignments of SARP regulators from different *Streptomyces* strains, highlighting the residues interacting with β (blue), σ R4 (cyan) and αCTD (purple). The black boxes highlight the positions conserved.

The downstream protomer also interacts with the RNAP β and β’ subunits. The ODB domain contacts the β-flap tip helix (FTH), the preceding loop (TPL) and the following loop (TFL) of the β flap by the helices α1 and β2, and the second antiparallel β sheet (Fig. 3D). The residues D71 and E76 of the ODB make salt bridges with the R805 of the β FTH. The R59 at the C-terminal of the ODB α1 forms a salt bridge with the E817 in TFL. The R62 and T63 of the strand β4 connecting the helices α1 and α2 contact the R816 in TFL and the K795 in TPL, respectively. The S106 in the loop connecting the strands β5 and β6 forms a hydrogen bond with the main chain oxygen atom of the K749 in the loop following the first strand of the β flap. The BTA domain forms interactions with the C-terminal of the β FTH via the N-terminal of the helix α3, of which the R173 side chain is parallel to TFL. The positively charged R67 and R69 region of β′ zinc-binding domain (ZBD) complements the negatively charged E76 and E77 of the HTH loop of the ODB domain (Fig. 3E).

To confirm the importance of the residues contacting RNAP subunits in transcription activation, they were replaced by alanine through site-directed mutagenesis. The mutants were purified and assayed in MangoIII-based *in vitro* transcription assays as the wild-type protein (Fig. 3F). Mutating residues R59, D71, E76 and R173 diminished the capability of SAS to active transcription, whereas no obvious difference was observed between the T63A, E77A, E246A, L247A and V249A mutants and the wild-type protein (Fig. S7). Mutating residues E176, L211 and L243 resulted in insoluble proteins when they were expressed under the same conditions as the wild-type protein. Some residues involved in RNAP interactions are highly conserved in SARP homologues across various *Streptomyces* strains (Fig. 3G), including T63 and D71 for interaction with the β subunit, L243, E246, L247, and V249 for interaction with σ R4.

### Interactions between SAS and αCTD

αCTD has versatility and important regulatory roles in modulating RNAP performance. In *E. coli* RNAP, it recognizes specific sequences called UP elements in some promoters (*33*). Many regulators have been reported to activate transcription by interacting with αCTD. The SAS-TIC structure demonstrates that both BTA domains interact with the αCTD, burying an interface area of ∼660 Å^2^ (Fig. 4A). The αCTD is positioned on top of the helix bundle of the downstream BTA domain. The helix α2 of αCTD interacts with the helix bundles of both upstream and downstream BTA domains and is almost perpendicular to them. The loops preceding and following helix α2, and the loop connecting helices α4 and α5 of αCTD make contacts with the helix bundle of the downstream and upstream BTA domain, respectively. The αCTD interface is predominantly positively charged (R259, R266, R287, K292), complementing the negatively charged interface of BTA domains (D245, E246 and D250 of the upstream BTA domain) (Fig. 4B). The residues R240 and L263 of the downstream BTA domain interact with αCTD by electrostatic and hydrophobic interactions respectively. As shown in Fig. 4C, the αCTD is required for SAS-dependent activation. Site-directed mutational studies further confirmed the importance of interfacial residues for SAS-dependent transcriptional activation. Moreover, most residues mentioned above for contacting the αCTD are highly conserved in SARP homologues, including R240, D245, E246, and L263.

**Fig. 4.**
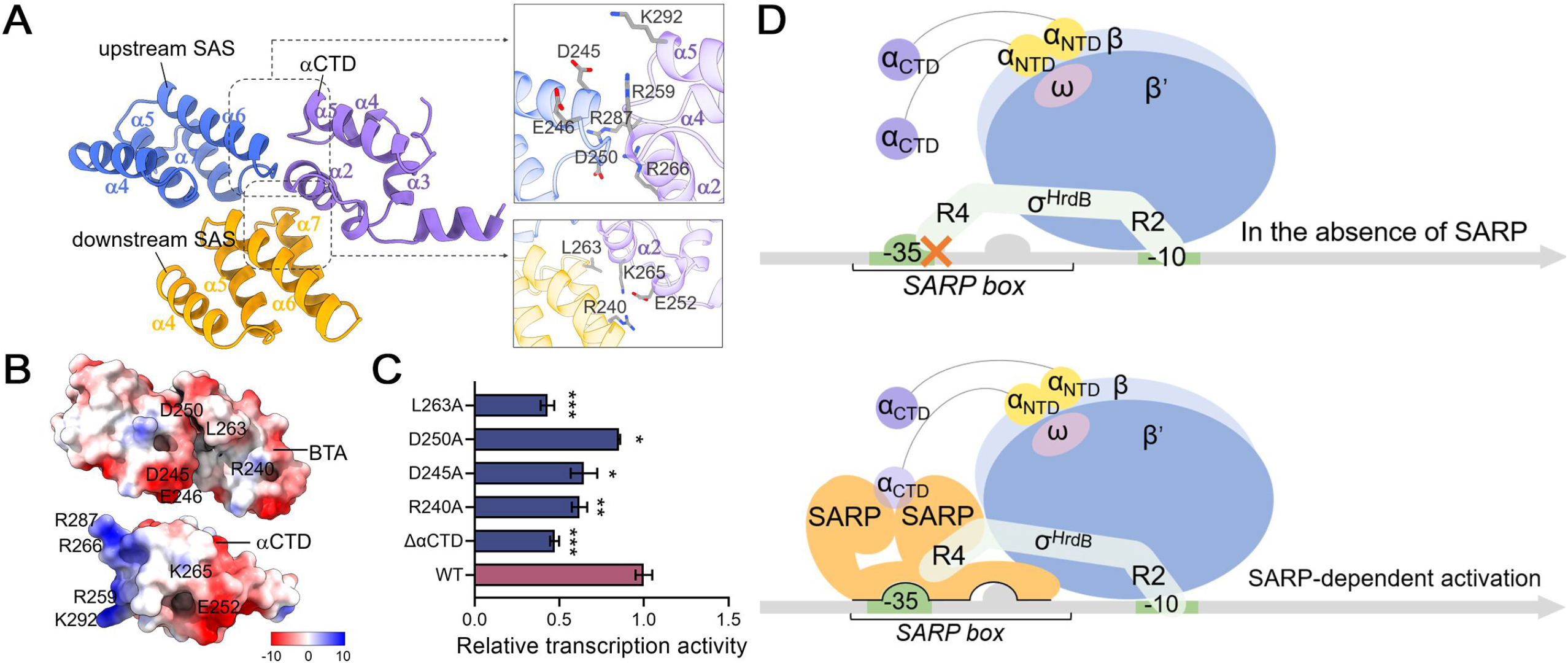
Interactions of SAS with RNAP αCTD. (A) BTA domains of both SAS protomers interact with the αCTD. Detailed interactions are shown in the grey box. (B) The electrostatic potential interface of BTA domains composed of R240, D245, E246, D250 and L263, and that of RNAP αCTD composed of E252, R259, K265, R266, R287 and K292. (C) Removing αCTD or mutating SAS residues involved in interactions with αCTD impaired transcription activation compared with wild-type SAS. Data are presented as mean ± SEM from three independent assays. ^*^*P* < 0.05; ^**^*P* < 0.01;^***^*P* < 0.001 in comparison with the wild-type SAS. (D) Proposed model for SARP-dependent transcription activation.

## 3. Discussion

Streptomycete promoters exhibit a wide diversity in sequences and transcription patterns to finetune the secondary metabolism to synthesize numerous natural products in response to environmental change. Genome-wide mapping of transcriptional start sites and ChIP-seq experiments of *S. coelicolor*, the model antibiotic-producing streptomycete, reveal that most promoters have a conserved −10 element, but a highly variable −35 element(*17, 18*). The promoter −35 elements of *M. tuberculosis*, another GC-rich actinomycete, are also lowly conserved(*34*). Guanosines at the position −13 and −14 and a transcriptional cofactor RbpA are shown to be important for promoters lacking a consensus −35 element(*35*). RbpA improves the interaction between the −10 element and σ R2 by binding to RNAP subunits and the promoter −13 and −14 positions(*32*).

SARPs are well-known transcription activators of antibiotic biosynthesis. Here, we report a strategy used by SAS to circumvent the recognition of a suboptimal −35 element and activate transcription. Given the comparable protein architecture of SARPs and their binding to similar motifs, SARPs are likely to function in a similar mode (Fig. 4D). The position of *afs box* (centered at −29.5) is different from the binding boxes of typical class II transcriptional activators (centered at −41.5). Consequently, the downstream SAS protomer is closer to RNAP and makes extensive contacts with β and β’ subunits (Fig. 3A). Within the scope of our investigation, SARP demonstrates the most comprehensive interaction with RNAP among the studied transcription activators, encompassing all subunits of RNAP except for ω, namely α, β, β’, and σ subunits. Particularly, SAS functions as an adaptor that physically bonds the −35 region and σ R4 together to compensate for the suboptimal −35 element. In reported class II activator-TIC structures (*22*), the σ R4 binds in the major groove of the −35 element and interacts with class II activators such as CAP (*21*), SoxS (*36*) and Rob (*37*) (Fig. S8). The transcriptional activation mode of SAS mostly resembles that of the mycobacterial transcriptional activator PafBC termed “sigma adaptation”(*30*) (Fig. S8F). PafBC inserts between PafBC-specific -26 element and σ R4 to facilitate the recognition of promoters with suboptimal −35 elements. Our study showed that the “sigma adaption” principle was widely adopted in actinobacteria, as indicated by the ubiquitous presence of SARP regulators. In addition, both SAS protomers make contacts with the αCTD, enhancing the ability to activate transcription by recruiting RNAP. Notably, although many activators in reported TIC structures are dimer, they only use one protomer to contact αCTD. In class I CAP-TAC structures, the downstream protomer contacts the αCTD (*21*). In contrast, the upstream protomer contacts the αCTD in class II TAP-TAC structures (*22*). Given the extensive interaction of SAS with RNAP, we raise a hypothesis that SAS may also activate transcription by a “pre-recruitment” mechanism, as exemplified by SoxS (*36*). But *in vitro* binding assay of SAS and RNAP demonstrates that SAS is unable to form a stable complex with RNAP in the absence of DNA (Fig. S9).

The majority of filamentous Actinobacteria genera are known to possess SARP-type regulatory genes, with examples including 98% of *Streptomyces* and 100% of *Salinispora*, while the occurrence in *Mycobacterium* is relatively lower, at only 42% (*38*). “Small” SARP regulators exhibit higher prevalence than “large” ones (*8*). These “small” SARPs hold significant potential as tools for activating biosynthetic gene clusters, as evidenced by the overexpression of SARPs to enhance the yield of natural products and the heterologous expression of SARPs to effectively activate the expression of silent gene clusters (*8, 39-41*).

*Sc*AfsR contains extra ATPase and TPR domains at the C-terminal that are postulated to be responsible for regulating the functions of the N-terminal SARP domain (Fig. 1A)(*9*). *Sc*AfsR belongs to the signal transduction ATPases with numerous domains (STAND) family (*42*). STAND proteins are presumed to keep in a monomeric inactive state by arm-based NOD–sensor autoinhibitory interactions or inhibitor binding in the absence of cognate inducer molecules, resembling a folded hand (*43-46*). Activation involves a multistep process of inducer binding, autoinhibition release, nucleotide exchange and conformational changes that allow oligomerization and switch the protein to an active state (*47*). This suggests that the C-terminal domains of *Sc*AfsR may potentially serve a similar role in sensing and transmitting signals. However, the precise mechanism remains unclear and requires further investigation.

In summary, transcription activators of the SARP family have been exclusively found in actinomycetes and play important roles in regulating secondary metabolism. Our detailed structural analysis of SAS-TIC provides a molecular basis for understanding SARP-mediated antibiotic regulation in streptomycetes and offers potential optimizations for antibiotic production.

## 4. Materials and Methods

### Plasmids

The genes encoding SAS (residues from 1 to 270 of *Sc*AfsR), RbpA and CarD were cloned from the genomic DNA of *S. coelicolor* M145, and inserted into the pET28a via *Nde*I and *EcoR*I restriction sites to obtain pET28a-*SAS*, pET28a-*rbpA* and pET28a-*carD* respectively. Plasmids carrying SAS mutants were constructed using site-directed mutagenesis. Primers used are listed in Table S1.

### Protein Expression and Purification

The vector pET28a-*SAS*, pET28a-*rbpA* and pET28a-*carD* were transformed into BL21(DE3) respectively. Induction of expression was achieved by adding 0.3 mM isopropyl-β-D-thiogalactopyranoside (IPTG) and incubated for 12 h at 16°C and 220 rpm. The cells were then harvested and resuspended in buffer A (50 mM Tris, pH 8.0, 500 mM NaCl, 10% glycerol, 1 mM β-mercaptoethanol and 5 mM imidazole). After purification by nickel-NTA, the eluate was further loaded onto a size exclusion chromatography column (SEC) (Superdex 200, Cytiva) equilibrated with buffer B (20 mM Tris, pH 8.0, 150 mM NaCl, 5% glycerol) and the purified proteins were stored at −80 °C. Purification of *S. coelicolor* RNAP and σ^HrdB^ was carried out as described previously (*27*).

### *In vitro* transcription

The transcription activities are evaluated by MangoIII-based transcription assay as previously described (*27*). DNA fragments containing the *afsS* promoter (−50 to + 15) and the MangoIII sequence were used as transcription templates. Primers used are listed in Supplementary Table S1. A gradient concentration range of SAS (ranging from 62.5 nM to 1000 nM) was combined with 10 nM of promoter DNA at 30°C for 10 minutes, followed by supplementation of 100 nM of RNAP-σ^HrdB^ and 1 mM of NTPs in transcription buffer (20 mM Tris–HCl, pH 8.0, 100 mM KCl, 5 mM MgCl_2_, 1 mM DTT, 4 U RNaseIn, 1 μM TO1–PEG–biotin and 5% Glycerol). The reactions were incubated for 15 min at 30 °C and stopped by 0.5 ng/μL (final concentration) heparin. In the investigation of the impact of RbpA and CarD on the functionality of SAS and the transcription of *afsS* promoter, as well as subsequent experiments involving SAS mutants, RbpA and CarD were co-incubated within the transcriptional system as global transcription factors (*48-50*). 500 nM (final concentration) of SAS or its variants, RbpA and CarD were used. The reaction without NTP was used as blank. A multi-detection microplate reader (Tecan Spark®) was used to measure fluorescence intensities. The emission wavelength was 535 nm and the excitation wavelength was 510 nm.

### Fluorescence Polarization (FP)

A 32-bp *afs* box (−45 to −14) consisting of a 5′-conjugated FAM was synthesized, annealled and used as the DNA probe. The reaction mixture contained 10 nM of labelled dsDNA and 0-5 μm purified SAS in binding buffer (20 mM HEPES, pH 7.5, 100 mM NaCl, 5 mM MgCl_2_, 2 mM DTT, 5% glycerol and 0.1 μg/mL of poly(dI-dC)). After incubation at 30 °C for 20 min, the fluorescence polarization of the reaction mixture was detected by a multi-detection microplate reader (Tecan Spark®) with excitation wavelength of 485 nm and emission wavelength of 535 nm. Data from technical triplicates were fitted to a binding equation Y=B_max_*X/(*k*_D-ave_+X) to obtain the *k*_D-ave_, where Y is the ΔmP measured at a given protein concentration (X) and B_max_ is the maximum ΔmP of completely bound DNA.

### Assembly of SAS-TIC complex

To construct SAS-TIC, we used a DNA scaffold engineered from the *afsS* promoter carrying a 6-bp pre-melted transcription bubble. The DNA was prepared by annealing nt-strand (100 μM final) and t-strand (110 μM). For assembly of SAS-TIC, 32 μM SAS was preincubated with 5 μM *afsS* promoter for 10 min at 25°C. 4 μM RNAP core, 16 μM σ^HrdB^, 16 μM CarD, and 32 μM RbpA were incubated for 10 min at 25°C, then mixed with preincubated SAS-DNA and incubated for another 10 min at 25 °C in assembling buffer (20 mM Tris-HCl pH 8.0, 50 mM KCl, 5 mM MgCl_2_, and 3 mM DTT). We then loaded the complexes onto a Superose 6 10/300 GL column (GE Healthcare), eluted with assembling buffer, and used them directly for cryo-EM grid preparation.

### Cryo-EM data acquisition and processing

SAS-TIC samples were added to freshly glow-discharged Quantifoil R1.2/1.3 Au 300 mesh grids. In a Vitrobot (FEI, Inc.), grids were plunge-frozen in liquid ethane after being blotted for 2 s at 16°C with 100% chamber humidity. The grids were imaged using SerialEM on Titan Krios 300 kV microscopes with a K2 detector. The defocus ranged from −1.0 to −2.0 μm, and the magnification was ×130,000 in counting mode. 32 frames per movie were collected with a total dose of 40 e^–^/Å^2^. For SAS-TIC, 3741 movies were collected.

The SAS-TIC data was processed using CryoSPARC suite v3.3.1. After motion correction, patch CTF estimation, manual exposure curation, and template picker using selected 2D classes from blob picker, two rounds of 2D classification were conducted (*51*). For SAS-TIC, 603659 particles were picked and used for ab-initio reconstruction (four classes). The selected particles (193644 particles) were used for the second round of ab-initio reconstruction (three classes) and heterogeneous refinement. The second 3D class (95223 particles, 49.2%) was selected and refined. Then, local resolution estimation and local filtering were conducted to give final maps. By the process of particle subtraction and masked local refinements to reserve only the signal around the SAS region, a local map with improved quality was obtained (Fig. S3).

### Cryo-EM model building and refinement

The initial models of *S. coelicolor* RNAPσ^HrdB^, RbpA, and CarD were generated from PDB:7X75 (*27*), 5TW1 (*32*) and 4XLR (*31*), respectively. The initial atomic model of SAS was generated by AlphaFold (*52*). The model of the promoter DNA was built in Coot (*53*). The models of RNAP, SAS and DNA were fitted into the cryo-EM density maps using ChimeraX (*54*). The model was refined in Coot and Phenix with secondary structure, rotamer and Ramachandran restraints (*55*). The validation was performed by MolProbity (*56*). The Map versus Model FSCs was generated by Phenix (Fig. S3). The statistics of cryo-EM data processing and refinement were listed in Table S2.

## Acknowledgments

**General**: We thank Jing Liu, Xinqiu Guo and Mengyu Yan at the Instrument Analysis Center (IAC) of Shanghai Jiao Tong University for supporting the cryo-EM data collection.

## Author Contributions

J.Z., S.L. and Z.D. designed the experiments and analyzed data.

Y.W., X.Y. and F.Y. purified the proteins and assembled the complex.

Y.W. performed *in vivo* biochemical assays.

Y.W., and X.Y. conducted cryo-EM data collection, image processing, atomic model, building and refinement.

J.Z. and Y.W. performed structural analyses and wrote the manuscript.

## Data Availability

Atomic coordinate has been deposited in PDB with accession numbers 8HVR (SAS-TIC). Two cryo-EM density maps have been deposited in Electron Microscopy Data Bank with accession number EMD-35047 (SAS-TIC), and EMD-35046 (αCTD-SAS-DNA from SAS-TIC).

## Supplementary Materials

Fig. S1. Purification of proteins.

Fig. S2. *In vitro* assays of the *Streptomyces coelicolor* AfsR SARP (SAS).

Fig. S3. Single-particle cryo-EM analysis of SAS-TIC.

Fig. S4. Comparisons of RNAP.

Fig. S5. Comparison of SAS with EmbR comprising an additional C-terminal forkhead-associated (FHA) domain.

Fig. S6. The binding of two SAS protomers in *afs box* results in a 20° bend of the helical axis of the dsDNA.

Fig. S7. *In vitro* assays of SAS mutants.

Fig. S8. Interactions of regulators with σ R4 and −35 element.

Fig. S9. SAS-RNAP complex was not formed in the absence of DNA.

Table S1. List of primer sequences used in this study.

Table S2. Cryo-EM data collection, refinement, and validation statistics.

